# GatorST: A Versatile Contrastive Meta-Learning Framework for Spatial Transcriptomic Data Analysis

**DOI:** 10.1101/2025.07.01.662625

**Authors:** Song Wang, Yuxi Liu, Zhenhao Zhang, Qin Ma, Qianqian Song, Jiang Bian

**Affiliations:** Biostatistics and Health Data Science, School of Medicine, Indiana University, Indianapolis, 46202, IN, USA; Department of Health Outcomes & Biomedical Informatics, College of Medicine, University of Florida, Gainesville, 32610, FL, USA; Department of Electrical and Computer Engineering, University of Virginia, Charlottesville, 22903, VA, USA; College of Life Sciences, Northwest A&F University, Yangling, 712100, Shaanxi, China; Department of Biomedical Informatics, College of Medicine, The Ohio State University, Columbus, 43210, OH, USA

**Author notes:** Corresponding author *Email address:*; (Jiang Bian). Equal contribution (Song Wang, Yuxi Liu, Zhenhao Zhang).

**Keywords:** Spatial transcriptomics, Graph Neural Networks, Contrastive learning, Meta learning, Spatial domain identification, Gene expression imputation, Batch effects removal

## Abstract

**Introduction:** Recent advances in spatial transcriptomics (ST) technologies have revolutionized our understanding of cellular functions by providing gene expression profiles with rich spatial context. Effectively learning spatial representations is crucial for downstream analyses and requires robust integration of spatial information with transcriptomic data. While existing methods have shown promise, they often fail to adequately capture both local (neighbor-level) and global (tissue-wide) spatial contexts. Moreover, they tend to rely heavily on augmentation strategies, which can introduce noise and instability.

**Objectives:** This study aims to introduce and demonstrate a novel, versatile framework called GatorST, which explicitly combines graph-based modeling with advanced learning strategies to generate spatially informed representations of ST data. GatorST is designed to improve various downstream tasks, including identification of spatial domains, gene expression imputation, batch effect removal, and trajectory inference.

**Methods:** GatorST constructs a spot-spot graph by connecting each node to its k nearest spatial neighbors and extracts two-hop neighborhood subgraphs to capture local context. At the global level, gene expression profiles are clustered using soft K-means to generate pseudo-labels, which serve as weak supervision signals within a contrastive learning framework. This process encourages the alignment of embeddings with shared pseudo-labels while separating those with different labels. GatorST further adopts an episodic training strategy inspired by meta-learning, wherein each episode consists of a support set for contrastive optimization and a disjoint query set for embedding classification, guided by the pseudo-labeled data. This design enables the model to classify unseen samples based on learned embeddings, thereby enhancing its generalization to new spatial contexts.

**Results:** Comprehensive comparisons with fifteen state-of-the-art methods across fourteen spatial transcriptomics datasets demonstrate that GatorST consistently achieves superior performance in identifying spatial domains, imputing gene expressions, and removing batch effects. The results showcase the versatility and strong generalization capabilities of GatorST across diverse tissue types and experimental settings.

**Conclusion:** GatorST effectively integrates spatial topology and global gene expression through graph-based modeling, pseudo-labeling, and contrastive meta-learning. This framework generates biologically meaningful representations and significantly improves key downstream tasks, including spatial domain identification, gene expression imputation, batch effect removal, and trajectory inference.

## 1. Introduction

Human organs and systems comprise a wide range of cell subpopulations, each contributing uniquely to physiological functions and processes. The spatial distribution of these cells and their interactions play a vital role in maintaining these functions [1]. Analyzing tissue regions and cells within their natural spatial context is therefore highly desirable. By examining the concordance and variability among tissue regions and cell types, researchers can gain a detailed understanding of intercellular communication, which has significant implications for uncovering disease mechanisms [2, 3].

Recent advances in spatial transcriptomics (ST) [4] provides gene expression profiles with spatial context, enabling unprecedented insights into cellular function, tissue organization, and microenvironmental interactions [5–11]. A key goal in ST analysis is to learn robust spatial representations that capture both gene expression and spatial architecture, as these representations are critical for downstream tasks such as identifying spatial domains, imputing gene expressions, removing batch effects, and inferring developmental trajectories [12]. However, achieving reliable and biologically meaningful representations is challenging due to the high dimensionality, sparsity, and technical noise inherent in ST data. Addressing these challenges requires computational methods that can effectively reduce dimensionality while preserving important spatial and biological signals.

Many recent methods have focused on clustering-based strategies to identify spatial domains, with representative approaches including BayesSpace [13], UTAG [14], SpaGCN [15], SpaceFlow [16], and BANKSY [17]. These methods leverage various strategies such as probabilistic modeling, integration of spatial graphs, and deep graph learning to capture local spatial patterns. Graph-based approaches (e.g., DeepST [18], CCST [19], STAGATE [20], Spatial-MGCN [21], GraphST [22]) have been especially promising, using graph neural networks to integrate gene expression and spatial relationships for improved clustering accuracy. Some methods incorporate selfsupervised contrastive learning to enhance representation robustness. Despite these advances, most existing methods inadequately capture both local (neighbor-level) and global (tissue-wide) contexts simultaneously. Additionally, reliance on simplistic corruption or heavy augmentations can introduce noise and instability, particularly in small-scale datasets, limiting the quality and interpretability of learned embeddings.

To overcome these limitations, we introduce GatorST, a novel framework tailored for spatial transcriptomics analysis. GatorST explicitly integrates local and global spatial contexts by constructing two-hop neighborhood subgraphs that preserve fine-grained spatial topology and generating global pseudo-labels via clustering of gene expression profiles as weak supervision. This design is incorporated into a contrastive learning framework that encourages spatial coherence and biological relevance in the learned embeddings. Furthermore, GatorST employs a meta-learning-inspired episodic training strategy to enhance generalization across diverse tissue types and spatial resolutions. Extensive evaluations across multiple datasets demonstrate that GatorST consistently outperforms state-of-the-art methods in spatial domain identification and gene expression imputation, offering a scalable, robust, and generalizable solution for spatial transcriptomics analysis.

## 2. Results

### 2.1. Overview of GatorST framework

GatorST is designed to learn robust and biologically meaningful representations of spatial transcriptomics data by integrating a graph-based approach with contrastive meta-learning. It begins by constructing a spot-spot graph, where each node represents a spatial spot characterized by its gene expression profile and spatial coordinates. Edges are defined using a top-k nearest neighbor strategy based on spatial proximity, enabling each node to capture local spatial context. To better model local structural relationships, a subgraph is extracted for each node, consisting of its two-hop neighbors, thereby enriching local representations and providing a more nuanced understanding of cellular microenvironments. Graph Convolutional Networks (GCNs) are then applied to these subgraphs to derive initial spot-level embeddings that integrate both gene expression and spatial structural information, serving as the basis for subsequent optimization.

Next, GatorST builds on these initial spot-level embeddings by employing a contrastive meta-learning approach to further refine and optimize them. Specifically, it first performs soft K-means clustering on the gene expression profiles to assign each spot a pseudo-label, providing weak supervision that promotes intra-cluster cohesion and inter-cluster separation. Leveraging these pseudo-labels, the meta-learning approach then applies an episodic training strategy, constructing meta-training tasks by sampling support and query sets from the pseudo-labeled data. In each episode, a two-step optimization process is conducted: first, contrastive learning aligns embeddings within the same pseudo-label group and separates those from different groups, explicitly making use of subgraph-based structural relationships. Second, a cross-entropy classification loss is applied to the query set to further fine-tune the embeddings and ensure adaptability to specific tasks. A combined objective function balances these two losses, resulting in final spot-level embeddings that are both structurally coherent and semantically informative.

Finally, the optimized spot-level embeddings demonstrate strong versatility and practical utility across various downstream tasks, including spatial domain identification, gene expression imputation, trajectory inference, and batch effect removal. This highlights GatorST’s potential as a flexible and powerful tool for comprehensive spatial transcriptomics analysis. Detailed implementation procedures are described in the Materials and Methods section.

### 2.2. GatorST outperforms benchmark methods in spatial domain identification

In this section, we aimed to show that GatorST could outperform benchmark methods in spatial domain identification. As illustrated in **Figure 2**, GatorST consistently outperforms existing benchmark methods in the five evaluation metrics: ARI, NMI, ACC, Purity, and Homogeneity. These results suggest the effectiveness and robustness of the GatorST in identifying spatially and biologically coherent domains. Specifically, GatorST shows strong performance across all slices of the human DLPFC dataset (slices #151507 to #151676), which are characterized by a complex and heterogeneous tissue architecture. On average, GatorST surpasses all competing state-of-the-art methods by a considerable margin. In particular, GatorST achieves an average improvement in ARI of more than 10% compared to the contrastive learning-based methods GraphST and CCST. This result suggests that GatorST more effectively captures spatial expression patterns, including spatial proximity and gene expression similarity, while preserving the underlying anatomical structure of the tissue. Importantly, the superiority of GatorST is not limited to human brain tissue. Significant performance improvements were observed in the human breast cancer and mouse brain anterior datasets. Across these diverse spatial contexts, GatorST consistently delivers superior performance across all evaluation metrics, demonstrating its generalizability and adaptability to varying spatial resolutions, gene expression profiles, and biological variability.

**Figure 1.**
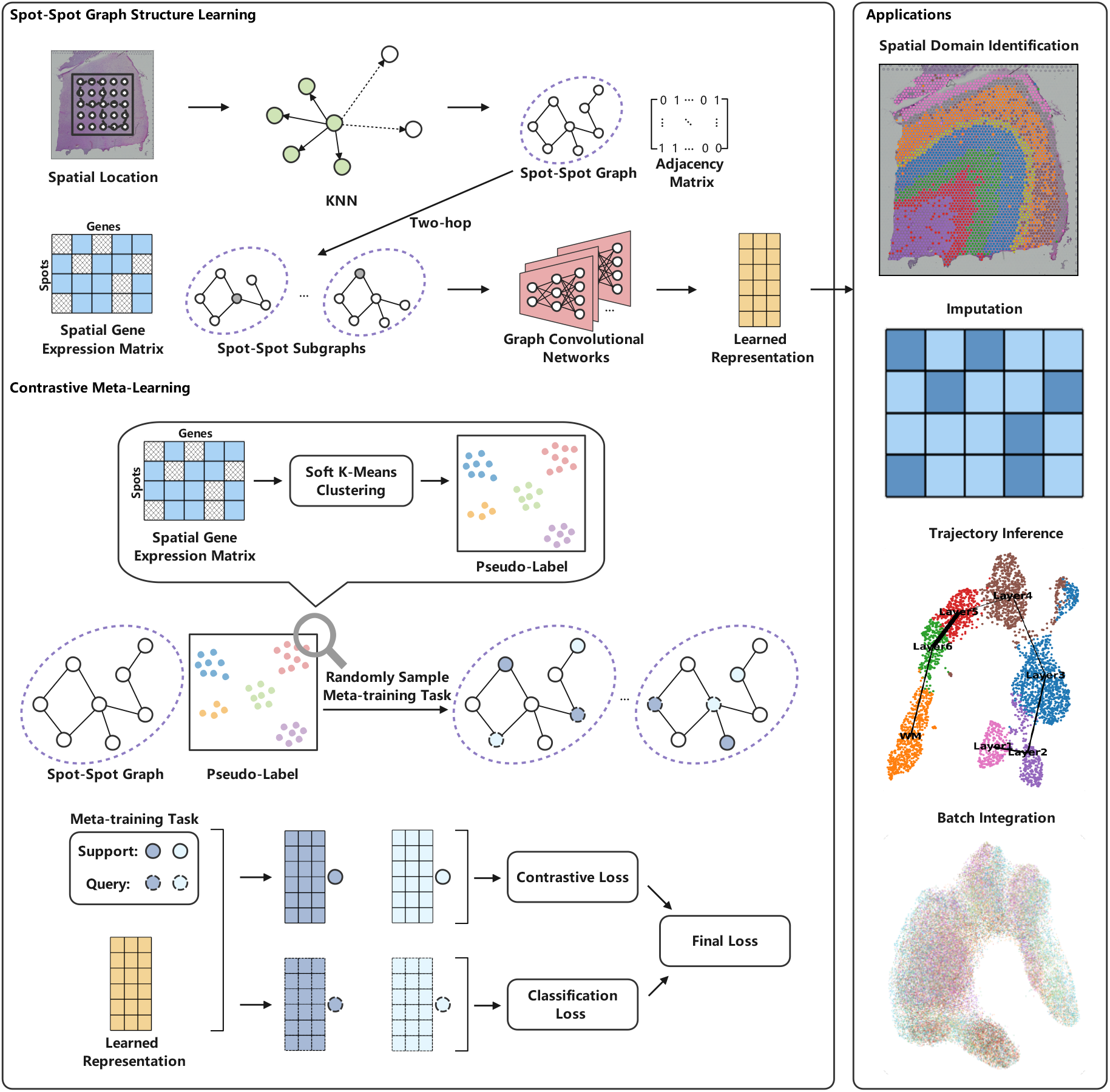
Overview of the GatorST framework.

**Figure 2.**
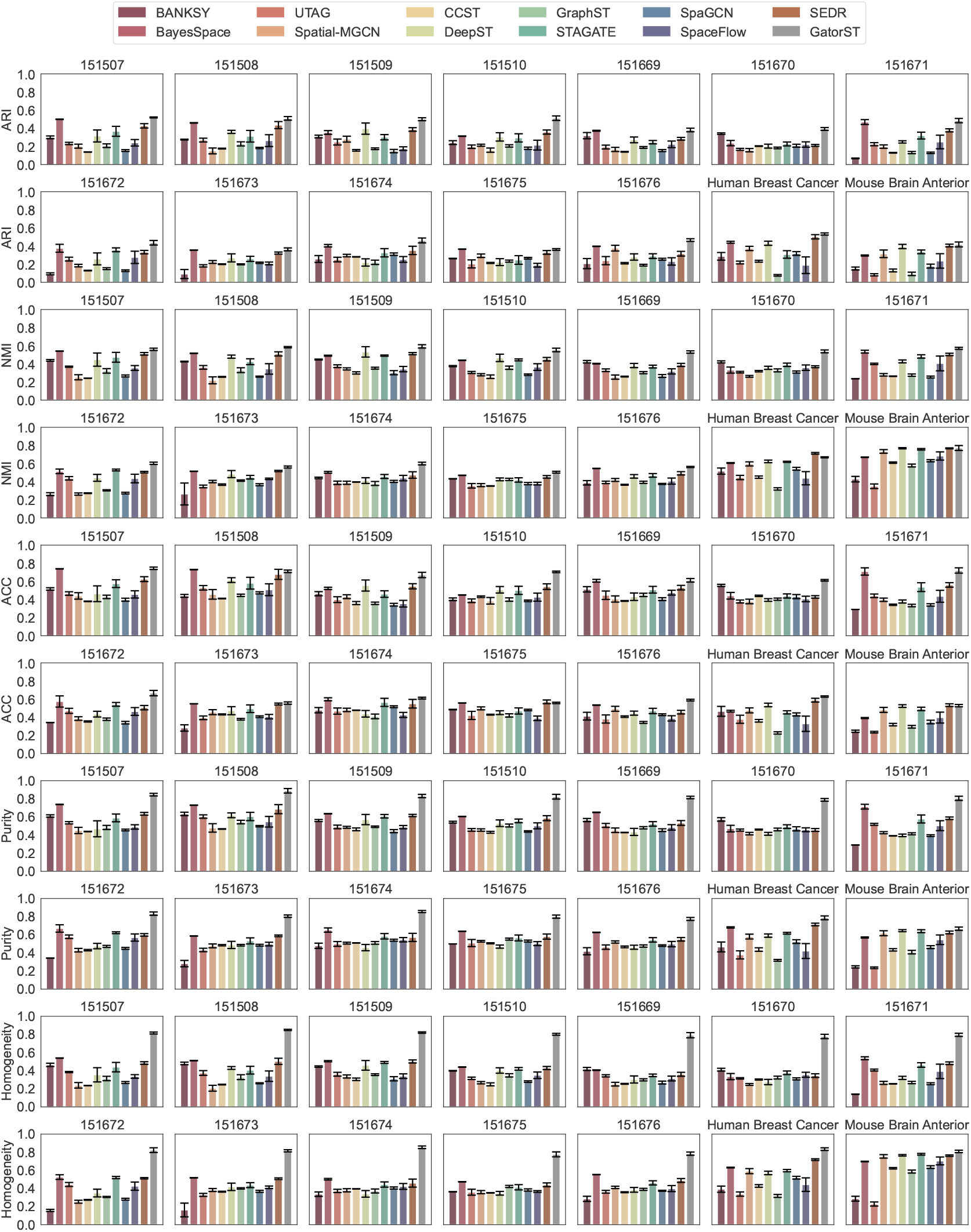
Clustering performance comparison using five clustering evaluation metrics ARI, NMI, ACC, purity, and Homogeneity on 14 spatial transcriptomic datasets. Higher values are better for all these metrics.

### 2.3. GatorST demonstrates superior imputation performance across diverse datasets

For gene expression imputation, we compared GatorST with a number of state-of-the-art spatial transcriptomics imputation methods, including SEDR [23], Spatial-MGCN, gimVI [24], and Tangram [25]. These methods can be classified into two groups: reference-based methods (i.e., gimVI and Tangram) and reference-free methods (i.e., SEDR and Spatial-MGCN). Specifically, gimVI and Tangram are designed to integrate spatial transcriptomics with single-cell RNA sequencing (scRNA-seq) data to accurately predict missing gene expression profiles. To ensure a fair comparison among all methods, we utilized their publicly available reference-free implementations, as these methods typically require matched scRNA-seq reference data, which may not always be accessible. In contrast, SEDR and Spatial-MGCN do not require matched scRNA-seq reference data. As shown in **Figure 3**, GatorST consistently achieved the highest PCC and the lowest L1 and RMSE across the majority of samples, demonstrating strong performance across diverse ST datasets. Notably, SEDR outperformed GatorST in terms of RMSE for the human breast cancer and mouse brain anterior datasets, suggesting that SEDR may have strengths specific to certain datasets. Overall, these results highlight the effectiveness and robustness of GatorST in spatial gene expression imputation tasks, particularly in reference-free contexts.

**Figure 3.**
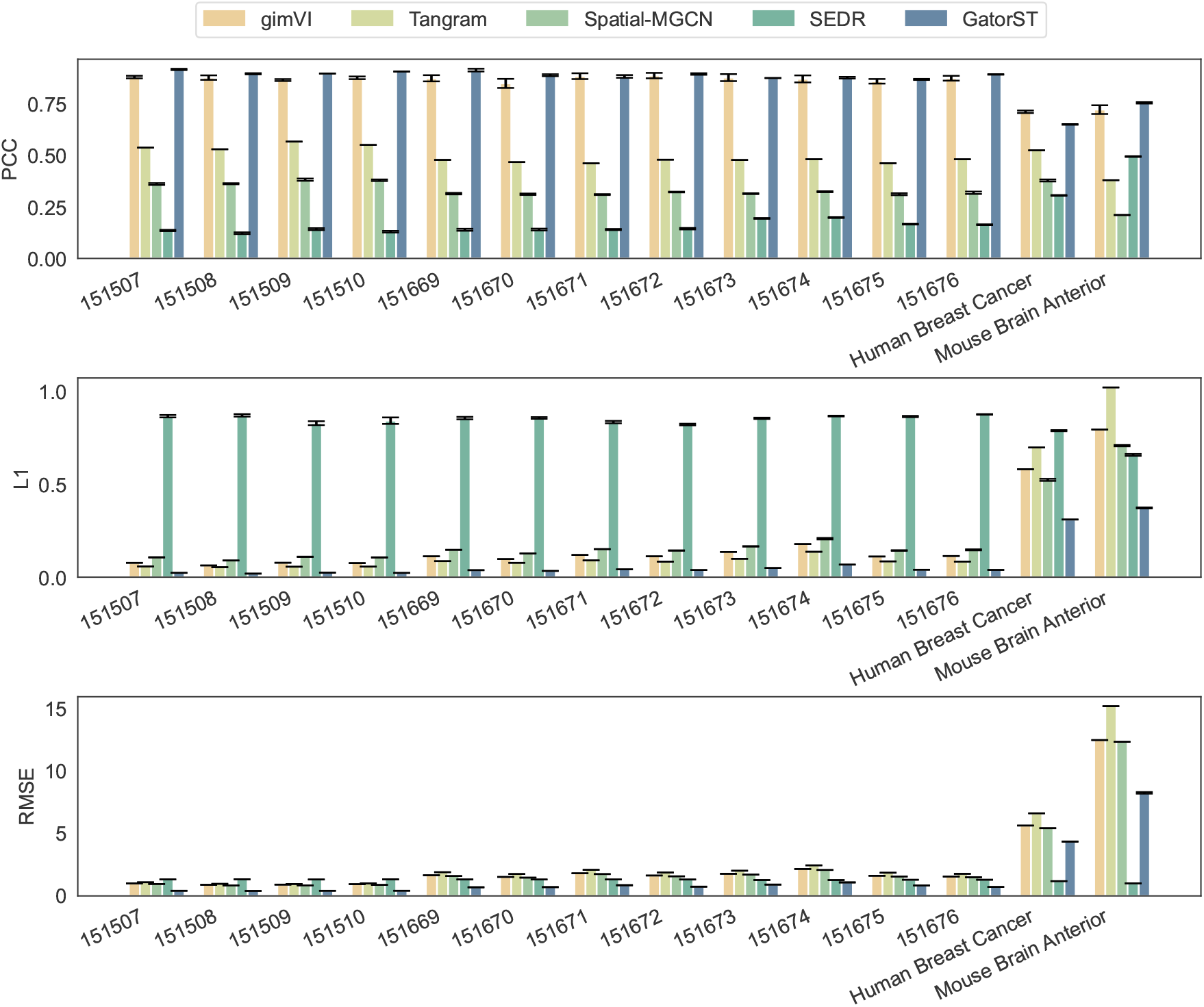
Imputation performance comparison on 14 spatial transcriptomic datasets using three evaluation metrics: PCC, L1 loss, and RMSE. A higher PCC value suggests better performance, while lower L1 loss and RMSE values suggest better performance.

### 2.4. GatorST accurately reveals anatomic layers in human prefrontal cortex

The human DLPFC dataset offers high-resolution spatially resolved transcriptomic profiles from 12 tissue slices, each capturing the structure of the human DLPFC. These slices include either four or six well-defined cortical layers, in addition to the underlying white matter (WM). The availability of detailed annotations makes this dataset a robust benchmark for evaluating the accuracy of spatial domain identification methods in recovering biologically meaningful cortical architecture. To assess the performance of the proposed GatorST and baseline methods, we applied all methods to four consecutive tissue slices from the dataset (slices #151673 to #151676). As shown in **Figure 4**, GatorST consistently achieved the highest ARI scores across all four slices, with values of 0.661, 0.662, 0.681, and 0.688, respectively. This consistent superiority demonstrates GatorST’s ability to effectively delineate cortical layer boundaries across varied tissue architectures. Among the competing methods, SEDR also showed competitive performance, achieving ARI scores of 0.585 on slice #151673 and 0.525 on slice #151675. However, its accuracy varied more significantly across different slices, suggesting a greater sensitivity to spatial noise or sample-specific variability. Collectively, these results highlight the superiority of GatorST in spatial domain identification, both in terms of overall accuracy and its adaptability to the structures of cortical tissue.

**Figure 4.**
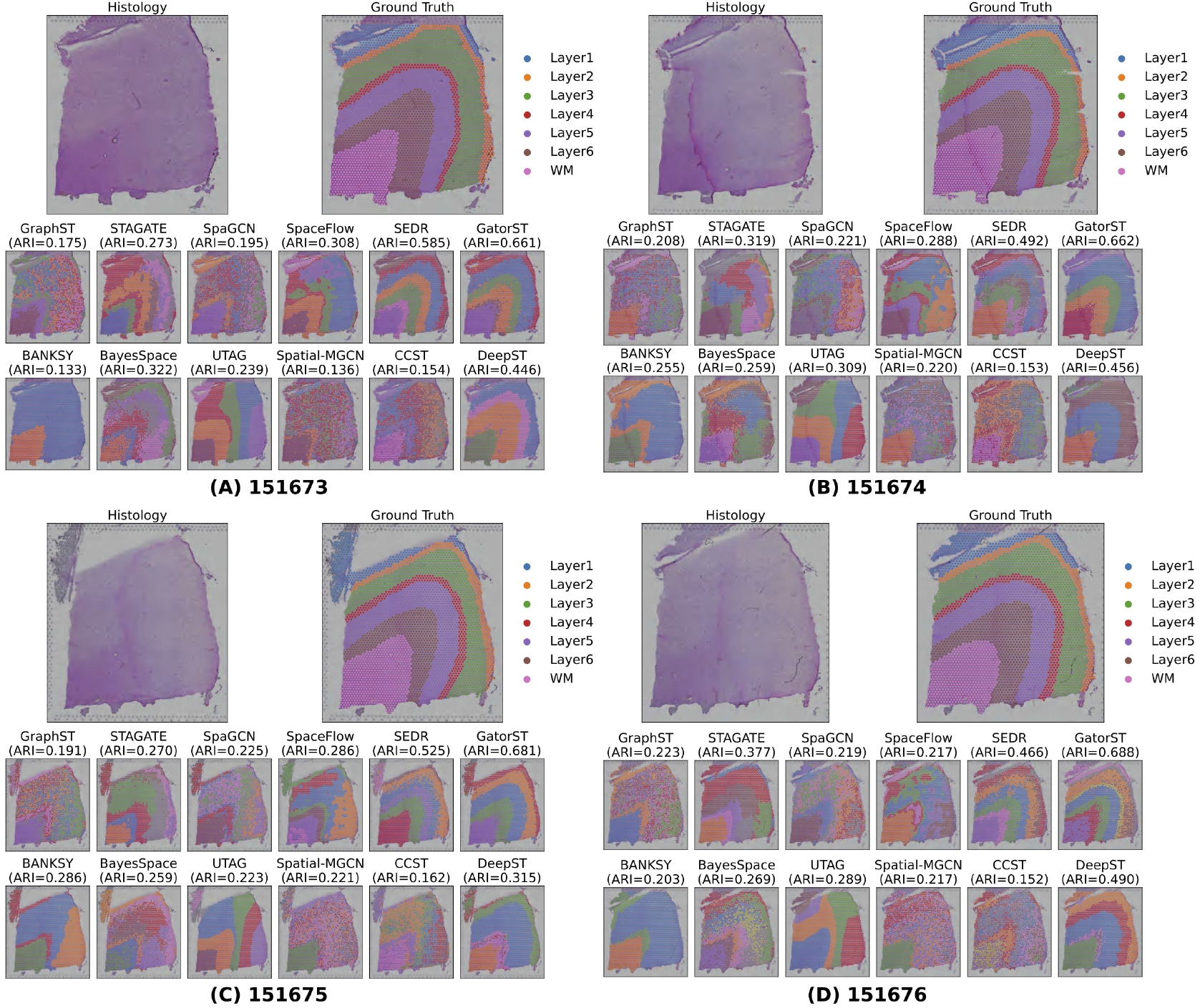
Comparison of manual annotations and clustering outputs for the representative slices #151673 to #151676 from the human DLPFC dataset.

### 2.5. Trajectory inference using spatial representations

To comprehensively evaluate the quality of the learned representations, we applied the proposed GatorST framework to two representative slices (#151675 and #151676) of the human DLPFC dataset. We visualized the learned embeddings using Uniform Manifold Approximation and Projection (UMAP) [26] and compared the outputs with those generated by several state-of-the-art baseline methods. As illustrated in **Figure 5**, GatorST produces a well-organized, sequential development of the cortical layers in slices 151675 and 151676 (Panels A and B). In the UMAP visualizations, GatorST displays distinct boundaries and minimal intermixing between adjacent cortical layers, highlighting its ability to preserve anatomical structure in the latent space. Each cortical layer forms a spatially coherent and distinct cluster, with Layer 1 and WM accurately localized at the respective cortical margins. Compared to baseline methods, GatorST reveals a more coherent laminar hierarchy characterized by sharper boundaries, less overlap, and improved separation between layers.

**Figure 5.**
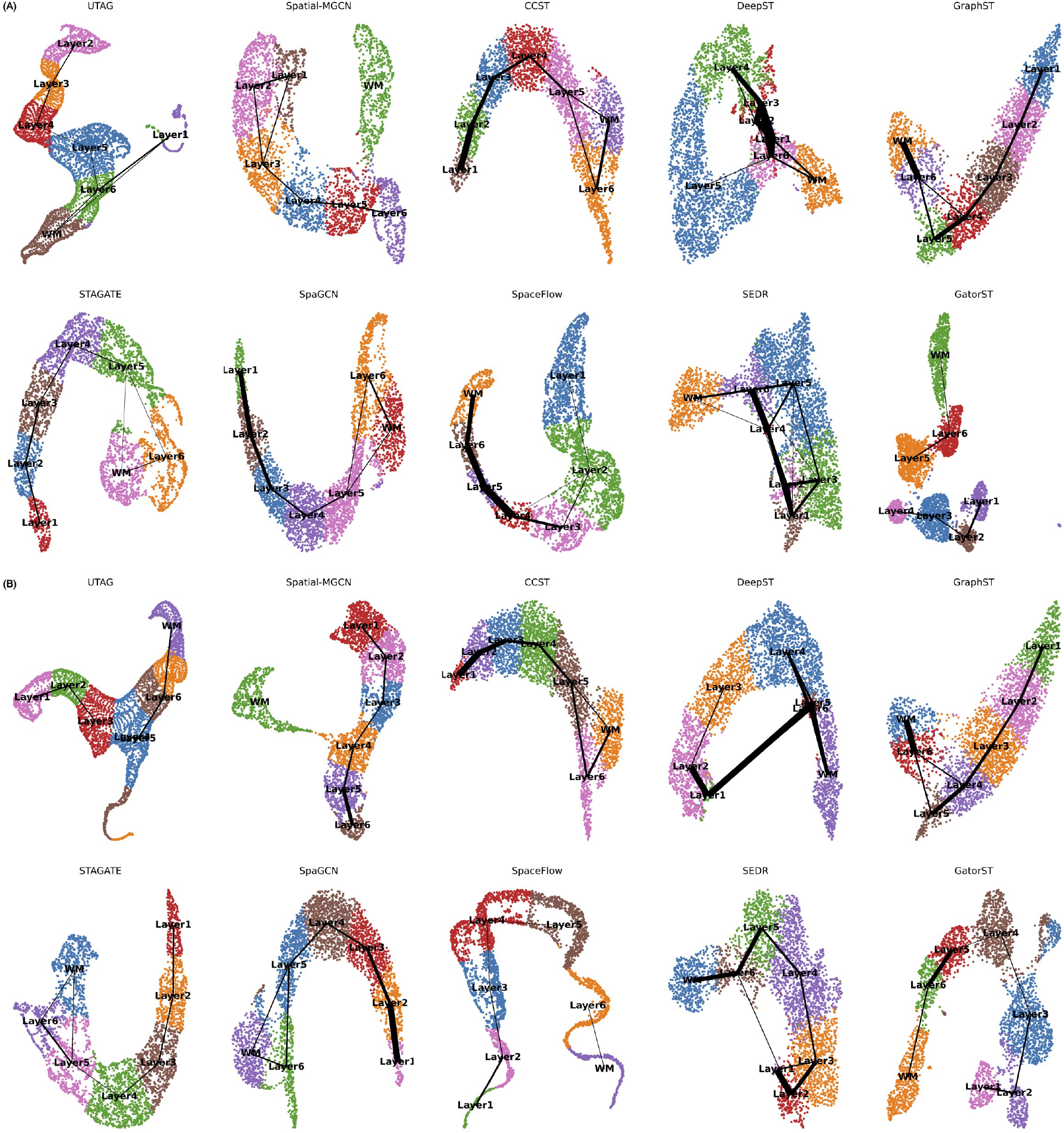
The UMAP and PAGA results of GatorST and baseline methods on human DLPFC slices #151675 and #151676.

To further evaluate the topological consistency of the latent space, we employed the partition-based graph abstraction (PAGA) [27], which constructs a graph-based representation of cluster connectivity based on transcriptomic similarity. In a PAGA graph, nodes represent clusters that correspond to cortical layers, while edges indicate the confidence of transcriptional transitions between clusters. Edge weights quantify the degree of transcriptomic continuity, with stronger connections reflecting gradual biological transitions and weaker or absent connections representing transcriptional boundaries. The PAGA graphs derived from GatorST embeddings reveal a linear, biologically plausible trajectory that reflects the expected spatial progression across cortical layers. This coherent structure contrasts with the PAGA graphs generated by baseline methods, which display fragmented topologies, spurious inter-cluster connections, or a lack of directional continuity. The well-defined and interpretable trajectories captured by GatorST further validate its ability to learn biologically meaningful gene expression patterns, reflecting the underlying cytoarchitecture of the human DLPFC.

### 2.6. GatorST effectively corrects for batch effects

To address the challenge of batch effects in spatial transcriptomics, we evaluated the use of joint embeddings across multiple batches by projecting them into a shared latent space. We benchmarked ten state-of-the-art methods, including our proposed GatorST, using the human DLPFC dataset to evaluate their effectiveness in preserving biological structure while integrating batch-specific variations, as illustrated in **Figure 6**. We utilized Harmony [28] as the batch integration algorithm because of its strong performance in single-cell data integration [29]. To quantify the performance of batch effect correction, we employed two metrics: cell type LISI (cLISI), which assesses the preservation of biological distinctions such as cortical layers, and the integration LISI (iLISI), which measures the degree of batch integration [28]. Higher iLISI values indicate better integration across batches, while lower cLISI values reflect superior preservation of biological structure.

**Figure 6.**
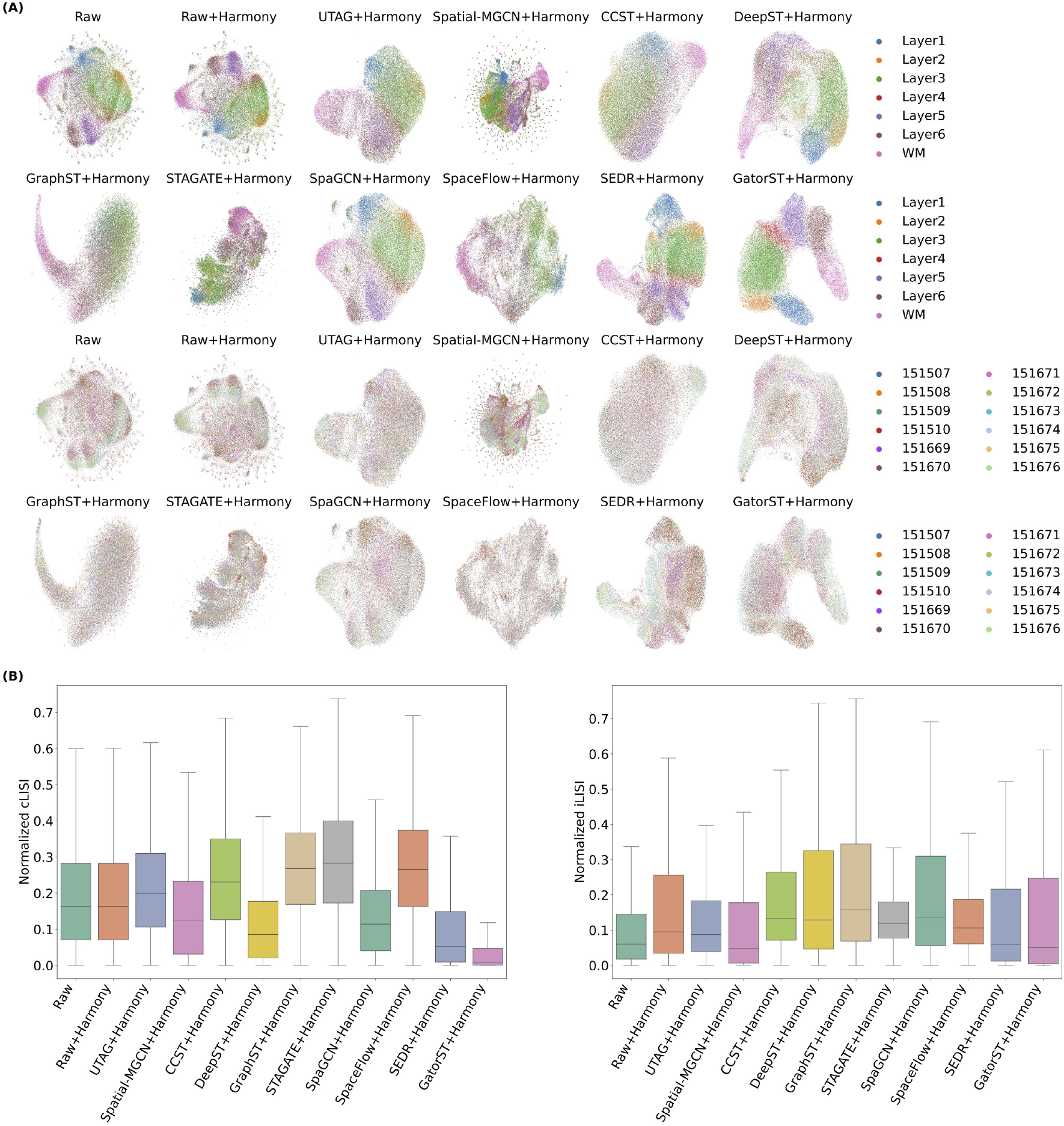
(A) Batch integration results for the human DLPFC dataset across nine baselines and the proposed GatorST method integrated with Harmony. (B) Quantitative assessment of integration performance was conducted using iLISI and cLISI scores. Higher iLISI values indicate better batch integration, while lower cLISI values reflect superior preservation of biological structure.

Specifically, we examined the Raw and Raw+Harmony to assess the impact of Harmony integration on Principal Component Analysis (PCA)-based embeddings. “Raw” refers to direct dimensionality reduction via PCA without any batch correction, while “Raw+Harmony” applies Harmony to the PCA-reduced representations. In **Figure 6B**, there was minimal difference in cLISI scores between the Raw and Raw+Harmony. However, Raw+Harmony demonstrated improved iLISI scores, suggesting enhanced batch integration compared to the uncorrected Raw. Among the baseline methods, STAGATE+Harmony achieved relatively high iLISI scores, indicating effective batch integration. Nonetheless, this improvement came at the expense of blurring biological structures, resulting in embeddings that failed to separate cortical layers and led to spatial ambiguity, as shown in **Figure 6A**. In contrast, SEDR+Harmony and GatorST+Harmony yielded well-separated and biologically coherent representations of cortical layers, effectively preserving not only layer-specific identities but also their developmental progression. While GatorST+Harmony achieved the lowest cLISI across all methods, indicating superior preservation of biologically coherent spatial architecture, its iLISI score was comparatively lower, reflecting a more conservative approach to batch integration. This outcome may be attributed to GatorST’s focus on local neighborhood structure and contrastive alignment based on pseudo-labels, which enhance intra-class cohesion but may resist excessive integration across batches. Overall, GatorST offers a structure-preserving approach that prioritizes biological fidelity over aggressive batch integration.

### 2.7. Parameter sensitivity analysis of the proposed GatorST framework

To further assess the effectiveness of GatorST under varying training conditions, we conducted a comprehensive parameter sensitivity analysis focusing on three critical hyperparameters: the learning rate, the batch size, and the temperature parameter *τ* used in the contrastive learning objective. In particular, the learning rate plays a key role in the convergence speed and overall optimization. We evaluated clustering performance across a broad range of learning rates from 10^*−*5^ to 10^*−*1^. As shown in **Figure 7**, performance was highly sensitive to this parameter. In particular, clustering performance improved with increasing learning rates up to a threshold, after which further increases led to performance degradation. Next we varied the batch size from 5 to 200 to examine its impact on clustering performance.

**Figure 7.**
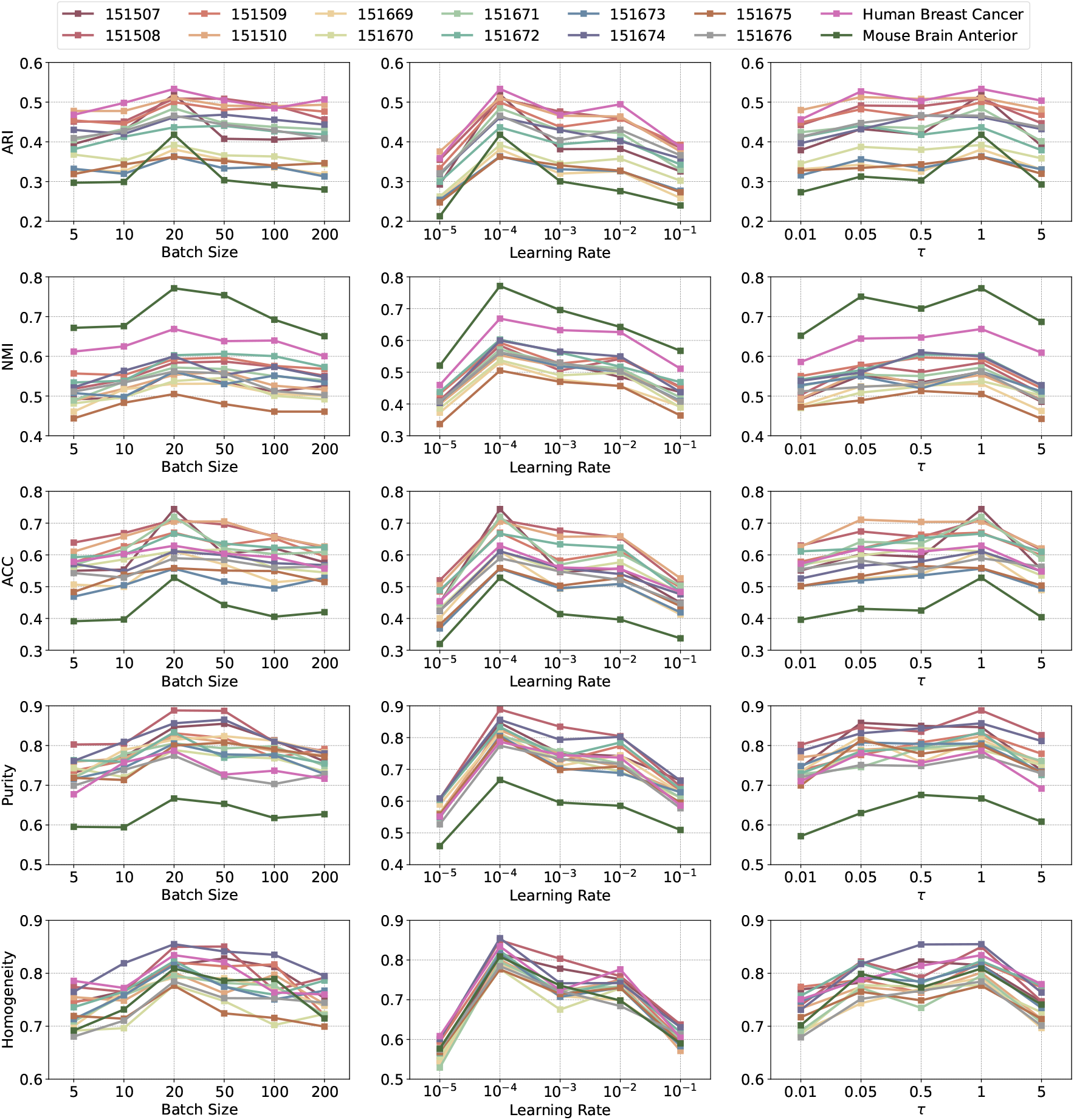
Hyperparameter analysis results using five clustering evaluation metrics: ARI, NMI, ACC, purity, and Homogeneity on 14 spatial transcriptomic datasets. The learning rate and the batch size, as well as the temperature parameter *τ* used in the contrastive learning objective were carried out in the analysis. Higher values are better for all these metrics.

The results indicated that intermediate batch sizes, particularly in the range of 20 to 50, consistently yielded higher ARI and NMI scores, providing an effective balance between clustering performance and training stability. The temperature parameter *τ* is a critical parameter in the contrastive learning objective, as it controls the sharpness of the similarity distribution and influences the separation of positive and negative sample pairs within the embedding space. We tested various values of *τ*, including 0.01, 0.05, 0.5, 1, and 5. Looking at **Figure 7**, moderate values of *τ* (e.g., 0.5 ∼ 1) achieved optimal performance across all evaluation metrics. Lower values such as *τ* = 0.01 excessively amplified contrastive penalties, leading to suboptimal performance, while higher values such as *τ* = 5 overly smoothed the similarity distribution, reducing the GatorST’s ability to discriminate between representations. Overall, the results of the parameter sensitivity analysis indicate that careful selection of the learning rate, batch size, and temperature parameter is critically important for improving the clustering performance of GatorST. Most importantly, GatorST maintains adaptability across a broad range of hyperparameter settings, highlighting its practical utility in ST data analysis.

### 2.8. Ablation studies of the proposed GatorST framework

To evaluate the contributions of the core components within the GatorST framework, we conducted extensive ablation studies across all benchmark datasets and clustering evaluation metrics. Specifically, we studied the following three variants: (i) GatorST_*α*_, where the contrastive learning objective was omitted; (ii) GatorST_*β*_, where the subgraph extraction was excluded; (iii) GatorST_*γ*_, where the Graph Convolutional Network (GCN) [30] was replaced with a Graph Attention Network (GAT) [31].

The results of the ablation study are shown in **Figure 8**. Among the three variants, GatorST_*α*_ displayed the most significant and consistent decline in clustering performance across the five metrics used. On average, the ARI decreased by more than 10%, with particularly significant declines observed in DLPFC slices 151507 and 151671, as well as in the mouse brain anterior dataset. Similarly, both NMI and Homogeneity showed approximately a 10% decrease. These results emphasize the crucial role of contrastive learning in aligning intra-class embeddings and maximizing inter-class distinction.

**Figure 8.**
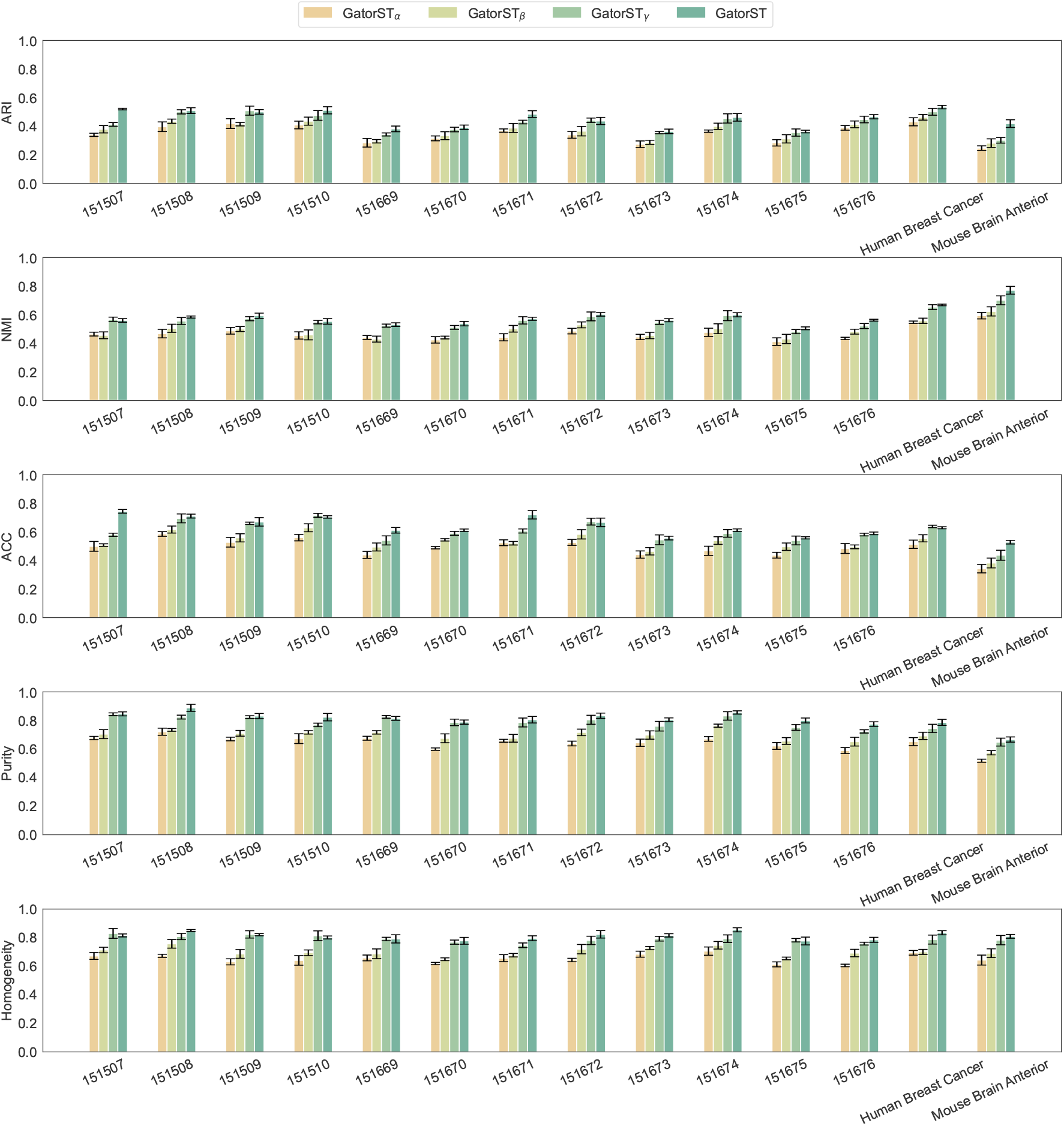
Clustering performance comparison across 14 spatial transcriptomic datasets between GatorST and its three variants.

GatorST_*β*_ demonstrated consistent yet moderate performance declines across datasets. While the decline in ARI was less severe compared to GatorST_*α*_, significant declines were observed in NMI, ACC, Purity, and Homogeneity, particularly in DLPFC slices 151510 and 151671, as well as in the mouse brain anterior dataset. For instance, both NMI and Homogeneity decreased by approximately 10%, highlighting the importance of incorporating localized structural context through two-hop subgraph extraction. These results suggest that subgraph-based modeling is vital for capturing microenvironmental relationships and preserving the spatial coherence of learned domains.

GatorST_*γ*_ yielded mixed outcomes. In certain DLPFC slices, like 151669, minor improvements in Purity and Homogeneity were observed. However, these improvements were inconsistent across datasets, with declines in Purity and Homogeneity observed in both the human breast cancer and mouse brain anterior datasets. ARI and NMI showed minor variations compared to GatorST for most datasets. While the GAT may introduce benefits in specific spatial contexts through its attention mechanisms [32], it does not consistently deliver optimal clustering performance. In contrast, the GCN provides a more balanced and efficient approach for modeling spatial transcriptomic data within the GatorST framework.

The results of the ablation study highlight the importance of the core components of GatorST. Both contrastive learning (GatorST_*α*_) and subgraph-based modeling (GatorST_*β*_) are critical for achieving improved clustering performance. These findings support the design rationale of GatorST, which integrates the capture of spatial topology, gene expressions, and the learning of discriminative feature representations into a unified framework for spatial domain identification.

### 2.9. The trade-offs between performance and computational efficiency

We evaluated the runtime and memory usage of all baseline methods and our GatorST using three datasets: slice #151669 of the human DLPFC dataset, human breast cancer, and mouse brain tissue. As shown in **Figure 9**, GatorST consistently achieved the highest ARI scores across all datasets, with ARI values of 0.3998 for slice #151669, 0.5415 for human breast cancer, and 0.4327 for mouse brain anterior, demonstrating its superior clustering accuracy. In terms of computational efficiency, GatorST demonstrated competitive runtime performance for all datasets. Notably, its runtime was consistently shorter than that of Spatial-MGCN and CCST, both of which showed longer runtimes despite delivering lower clustering accuracy. While GatorST showed relatively higher memory usage compared to most baseline methods (typically ranging from 2^12^ to 2^13^ MB), it effectively utilized these additional computational resources to deliver improved clustering results. For instance, in the human breast cancer and mouse brain tissue datasets, GatorST, despite its larger memory footprint, outperformed SpaGCN and UTAG, which had lower resource demands but yielded suboptimal ARI scores. These results highlight a significant trade-off: although GatorST requires greater memory, it leverages that capacity to achieve stateof-the-art clustering performance while maintaining runtime efficiency. This balance between clustering accuracy and computational cost makes GatorST as a powerful tool for studying spatial domain identification. Nevertheless, it is important to acknowledge that GatorST’s high memory requirements may limit its applicability in resource-constrained environments.

**Figure 9.**
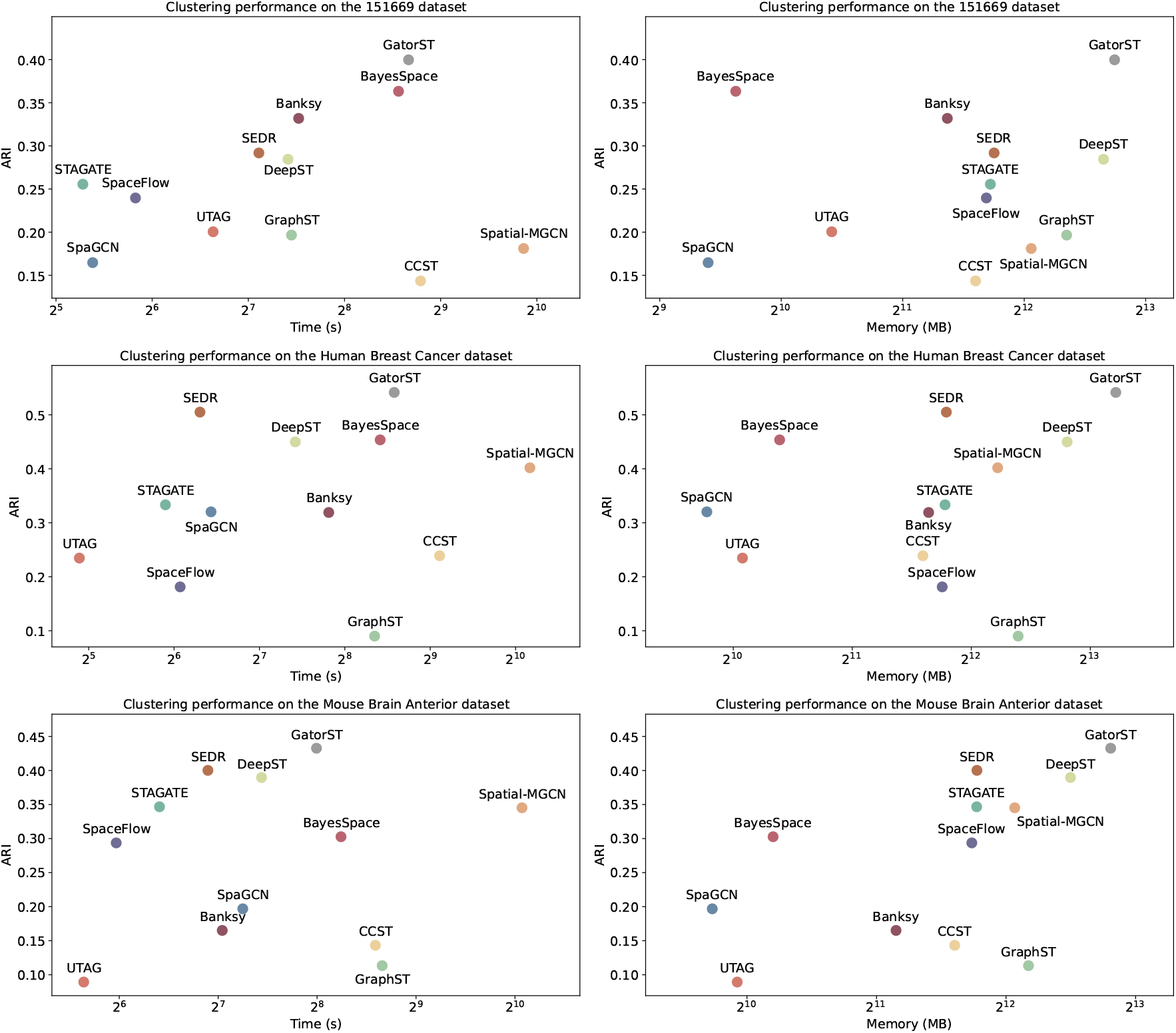
Clustering performance, runtime, and memory usage of baseline methods and our proposed GatorST across the slice #151669 of the human DLPFC dataset, human breast cancer, and mouse brain anterior tissue. Each scatter plot illustrates the trade-offs between clustering accuracy (measured by ARI) and computational efficiency.

## 3. Materials and Methods

### 3.1. Spot-Spot Graph Structure Learning

Let 𝒢 = (𝒱, ℰ, **X**) = (**A, X**) represent a spot-spot graph. In the graph, 𝒱 = {*v*_1_, *v*_2_, …, *v*_*C*_} represents the set of nodes/spots, where *C* is the total number of spots. *ℰ* represents the set of edges that connect the nodes. **X** ∈ R^*C*×*g*^ is the gene expression matrix obtained after preprocessing, and *g* is the total number of genes. The adjacency matrix **A** ∈ {0, 1}^*C*×*C*^ encodes the connectivity of the graph, where **A**_*ij*_ indicates the presence or absence of an edge between spots *i* and *j*.

To construct the spot-spot graph 𝒢, a similarity score for each pair of nodes is calculated and then used to define the entries in the adjacency matrix. The adjacency matrix **A** can be formalized as follows:

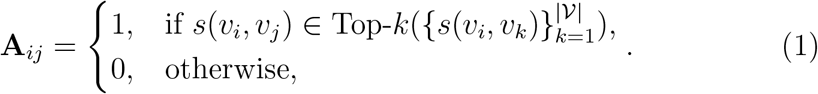

Up to *k* nearest neighbors (i.e., nodes with the highest similarity scores) are selected for each node, and edges are established to connect them. The similarity score *s*(*v*_*i*_, *v*_*j*_) between the nodes *v*_*i*_ and *v*_*j*_ is calculated using their spatial locations. The spatial location of a node *v*_*i*_ is represented by a location vector **z**_*i*_ ∈ ℝ^*z*^, where *z* represents the corresponding dimensionality. Accordingly, the similarity score between two nodes is calculated as follows:

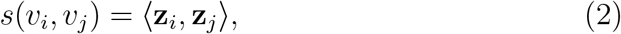

where **z**_*i*_ and **z**_*j*_ are the location vectors of *v*_*i*_ and *v*_*j*_, respectively.

Based on the foundation established by the spot-spot graph 𝒢, a subgraph is extracted for each node to represent its structural relationships. In particular, the subgraph for a given node consists of its two-hop neighboring nodes. We also tested incorporating 3-hop neighboring nodes; however, this did not improve performance and significantly increased computational costs. We take an example for node *v*_*i*_, and its subgraph 𝒢_*v*_ can be defined as follows:

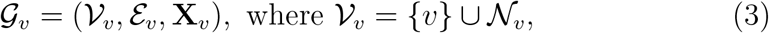

where 𝒩_*v*_ is the set of two-hop neighboring nodes of *v* can be further written as follows:

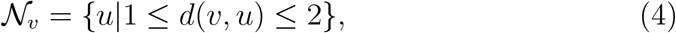

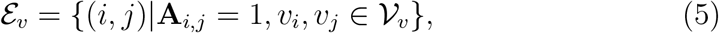

where the gene expression data for the nodes is represented by the matrix 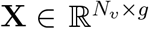. Here, *N*_*v*_ = |𝒱_*v*_| is the number of nodes and *g* is the dimensionality of the gene expression features. In order to learn the representation of the node *v*, a Graph Convolutional Network [30] is applied to its subgraph 𝒢_*v*_ as follows:

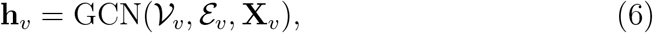

where 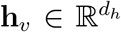 is the learned representation of node *v* and *d*_*h*_ the dimensionality of the representation. The learned node representations derived from subgraph-based graph convolution are subsequently utilized for various downstream tasks, such as spatial domain identification. To further enhance the quality and generalizability of these representations, we introduce a contrastive meta-learning framework, which integrates both local graph structure and global expression patterns into a unified optimization framework. The details of this framework are described in the following subsection.

### 3.2. Contrastive Meta-Learning

In this subsection, we present a contrastive meta-learning framework that leverages cluster-aware pseudo-labels as weak supervision to guide the alignment of spot embeddings. This design not only improves intra-cluster coherence and inter-cluster separation but also facilitates robust generalization across spatial contexts. In the following subsections, we detail the two key components of this framework: (i) the generation of pseudo-labels via soft Kmeans clustering, and (ii) an episodic training strategy that jointly optimizes contrastive and classification objectives.

#### 3.2.1. Cluster-Aware Pseudo-Labeling

Given the graph representation 𝒢= (𝒱, *ε*, **X**) = (**A, X**), each node *v ∈*𝒱is assigned a pseudo-label by applying the *K*-means clustering to the node feature matrix **X**. The goal of *K*-means clustering is to partition the nodes into *K* distinct clusters by minimizing the sum of squared distances between each node and its assigned cluster centroid.

Let {*µ*_1_, *µ*_2_, …, *µ*_*K*_} represent the set of *K* cluster centroids and ^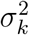^ represent the variance associated with cluster *k*. Each node *v*_*i*_ has an associated probability distribution over the clusters as follows:

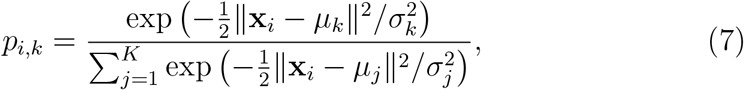

where *p*_*i,k*_ represents the probability that node *v*_*i*_ belongs to cluster *k*, ensuring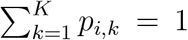. Instead of applying conventional hard assignments to cluster membership, the cluster centroids are updated based on the soft assignments derived from the probabilities:

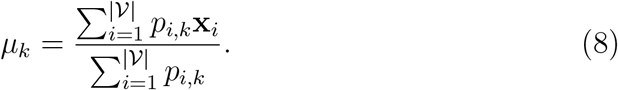

Accordingly, the clustering process consists of two main iterative steps: (i) computing *soft assignments p*_*i,k*_ and (ii) updating *centroid updates µ*_*k*_. These steps are repeated until convergence is reached. Finally, each node receives a *continuous pseudo-label* in the form of a cluster membership probability vector as follows:

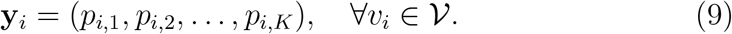

We further consider *y*_*i*_ as the hard-coded label of *v*_*i*_:

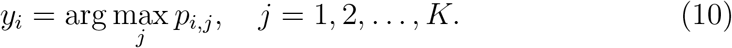

The resulting *y*_*i*_ serves as the pseudo-label for each node. Through this pseudo-label generation process, we establish a weak supervision signal based on soft clustering. These pseudo-labels are then utilized during meta-training to guide the joint optimization of node representations through both contrastive and classification objectives.

#### 3.2.2. Multi-objective optimization

With the pseudo-labels at hand, we are able to build an episodic training strategy that consists of multiple meta-training tasks. Meta-learning, often referred to as “learning to learn”, trains the model to efficiently adapt to new tasks by optimizing across a distribution of tasks rather than a single task [33–35]. This approach is particularly well suited for ST data analysis, as it enables the model to generalize across diverse spatial and biological contexts, addressing the inherent variability in sample distributions. To be specific, we begin by sampling a support set 𝒮 and a query set 𝒬 to establish a metatraining task. The support set 𝒮 consists of *N* randomly sampled classes from the *K* classes (i.e., clusters), with up to *M* nodes randomly selected from each of these *N* classes. Accordingly, 𝒮 is also known as a *N* -way, *M* - shot support set. In the same vein, the query set 𝒬consists of up to *Q* nodes from the same *N* classes, and these sampled nodes are distinct from those in *𝒮*. By doing so, each meta-training task can be established as follows:

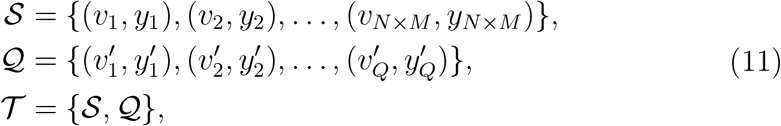

where *v*_*i*_ (or *q*_*i*_) represents a node in 𝒱 and *y*_*i*_ (or 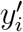) is the pseudolabel associated with *v*_*i*_ (or 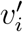).

During each episode, a two-step optimization process is implemented. First, contrastive learning is integrated with pseudo-labeling within each meta-training task 𝒯. Let 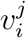 represent the *j*-th node in the *i*-th class within the support set *S*, with its learned representation represented as 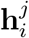, where *i* = 1, 2, …, *N* and *j* = 1, 2, …, *M*. Accordingly, the contrastive loss for the support set can be written as follows:

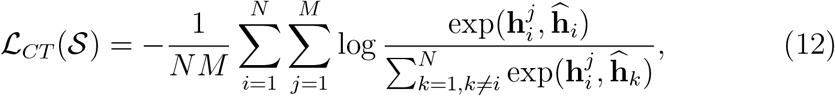

where 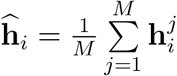 represents the representation of the *i*-th class, calculated by averaging the representations of *M* nodes that belong to the *i*-th class. By minimizing contrastive loss ℒ_*CT*_, we encourage the alignment of node embeddings that share the same pseudo-label while maximizing the separation between embeddings of nodes with different pseudo-labels.

Second, for the nodes in the query set, task-specific classification is carried out by assigning each node a label from the *N* available classes. This classification enables the fine-tuning of the learned representations, ensuring effective adaptation to the specific meta-training task 𝒯. Accordingly, the classification loss can be calculated as follows:

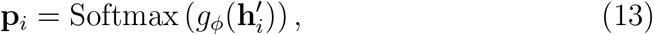

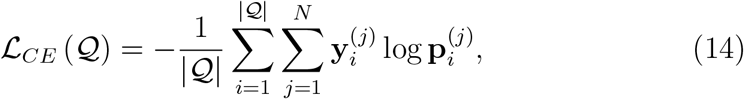

where **p**_*i*_ ∈ℝ^*N*^ represents the probability distribution over the *N* classes in the meta-training task 𝒯 for the *i*-th query node 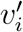 in 𝒬. A note of caution is due here 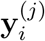 is the pseudo-label value with respect to the *j*-th class among the *N* classes. The value 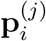 corresponds to the *j*-th component of **p**_*i*_.

By combining the contrastive loss *ℒ*_*CT*_ on *𝒮* and the classification loss ℒ_*CE*_ on 𝒬, the final optimization objective for the meta-training task can be formalized as follows:

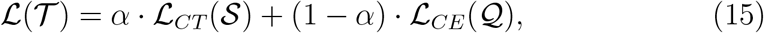

where *α ∈* [0, 1] is a hyperparameter that balances the contributions of both losses. Through the training process of contrastive meta-learning, we obtain node embeddings that are both structurally coherent and semantically discriminative. These representations can be directly applied to various spatial transcriptomics analysis tasks, including spatial domain identification, gene expression imputation, trajectory inference, and batch effect removal. In the following sections, we detail the implementation of each task.

### 3.3. Application tasks

#### 3.3.1. Spatial Domain Identification

Once the GatorST framework is properly trained, the learned spot representations can be utilized for the task of spatial domain identification. To achieve this, we adopt a clustering-based strategy that partitions all learned representations into distinct clusters, each corresponding to a spatial domain. Let 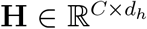 denote the matrix of learned spot representations, where *C* is the number of spots and *d*_*h*_ is the dimensionality of each representation.

Accordingly, we apply K-means clustering to **H** by optimizing the following objective function:

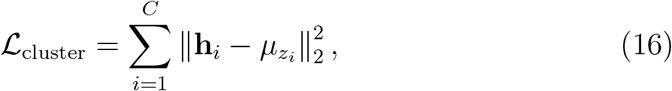

where 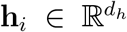is the representation of the *i*-th spot (i.e., the *i*-th row of **H**), *z*_*i*_ ∈ {1, …, *K*} is the cluster assignment of spot *i*, and 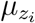 is the centroid of the *z*_*i*_-th cluster. This objective function is used to encourage each representation to be close to the centroid of its assigned group, effectively grouping spots with similar features into the same spatial domain.

#### 3.3.2. Gene Expression Imputation

GatorST also enables imputation of gene expression values by projecting the learned embeddings back into the gene expression space. A linear transformation is applied:

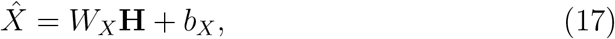

where *W*_*X*_ and *b*_*X*_ are learnable parameters. The imputed gene expression matrix 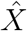 is then compared to the original expression matrix *X* using the Mean Absolute Error (MAE):

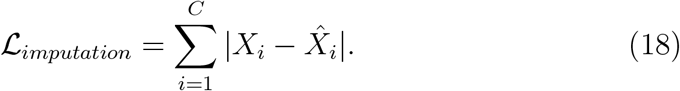

This objective guides the model to minimize deviations from the original measurements.

#### 3.3.3. Trajectory Inference

To reconstruct developmental or spatial trajectories, GatorST embeddings can be utilized as input for established trajectory inference tools such as PAGA [27]. Specifically, we applied the PAGA method to the embeddings derived from two representative DLPFC slices: #151675 and #151676. PAGA then constructs a connectivity graph where nodes correspond to discrete clusters (e.g., cortical layers), edges indicate transcriptional transitions between these clusters, and edge weights reflect the degree of transcriptomic continuity. This setup allows GatorST to support biologically plausible trajectory modeling. To enhance visualization, UMAP [26] is employed to project the learned embeddings into a two-dimensional space, providing a clear and intuitive overview for subsequent trajectory reconstruction.

#### 3.3.4. Batch Effect Removal

GatorST addresses batch effects by integrating Harmony into its framework. Harmony is applied to the latent embeddings across all 12 DLPFC tissue slices using the harmonypy (v0.0.9) implementation. The resulting batch-corrected embeddings are used for both visualization (i.e., UMAP) and quantitative assessment via LISI metrics. LISI metrics were computed using the *compute lisi* function provided in Harmonypy (v0.0.9), and two metrics were considered: cell-type LISI (cLISI) and integration LISI (iLISI) [28]. cLISI evaluates local cell-type diversity, where lower values indicate better preservation of biological structure. iLISI assesses mixing across datasets, with higher values reflecting superior batch integration. To facilitate comparison, both scores are normalized to the range [0, 1]:

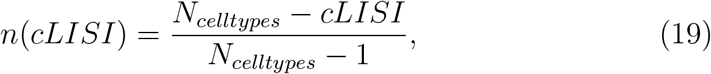

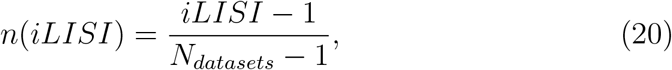

where *N*_*celltypes*_ is the number of clusters and *N*_*datasets*_ is the number of datasets.

### 3.4. Benchmark datasets

In this study, we assessed the performance of GatorST using various publicly available spatial transcriptomics datasets across different tissue types and spatial contexts. Specifically, we included the LIBD human dorsolateral prefrontal cortex (DLPFC) dataset [36], which consists of 12 tissue slices obtained using the 10x Visium platform (http://research.libd.org/spatialLIBD/). Additionally, we utilized datasets from the 10x Genomics Data Repository, including samples from human breast cancer (https://www.10xgenomics.com/datasets/human-breast-cancer-block-a-section-1-1-standard-1-1-0) and mouse brain tissue (https://www.10xgenomics.com/datasets/mouse-brain-serial-section-1-sagittal-anterior-1-standard-1-1-0). Annotations for these two datasets are accessible from the SEDR [23]. These datasets include a wide range of gene expression profiles and spatial resolutions, providing a comprehensive benchmark for assessing the generalizability and robustness of the proposed method. Notably, no pre-filtering was applied to the raw gene expression data. All available data were included in the analysis to ensure an unbiased and thorough evaluation.

### 3.5. Baseline methods

To thoroughly assess the clustering performance of GatorST, we compared it with various state-of-the-art spatial transcriptomics analysis methods. These included the Bayesian approach BayesSpace, graph-based models UTAG, Spatial-MGCN, DeepST, GraphST, STAGATE, SpaGCN, and SpaceFlow, as well as contrastive learning-based methods CCST and GraphST. Additionally, we compared GatorST with other representative methods, including BANKSY and spatially embedded deep representation (SEDR) [23]. BANKSY is a unified embedding framework that enhances gene expression profiles with features from spatial neighborhoods to jointly perform cell typing and domain segmentation. SEDR is a deep learning framework that integrates gene expression and spatial information by combining an autoencoder with a variational graph autoencoder.

### 3.6. Performance evaluation metrics

We conducted a rigorous evaluation of the proposed method using five commonly used metrics for clustering performance, including the Adjusted Rand Index (ARI), Normalized Mutual Information (NMI), clustering accuracy (ACC), purity, and Homogeneity. Specifically, ARI measures the agreement between the predicted and reference clusterings while accounting for randomness. This makes ARI particularly useful for assessing the reliability of clustering, particularly in the presence of overlapping classes. NMI quantifies the mutual dependence between the predicted and actual labels. By normalizing mutual information, it accounts for label distribution imbalances, with higher scores suggesting stronger agreement. ACC reflects the proportion of correctly assigned instances. It is calculated by optimally aligning the predicted clusters with the ground truth. Purity evaluates how well each predicted cluster predominantly consists of members from a single ground truth class. It reflects the coherence within each cluster. Homogeneity measures the degree to which clusters consist entirely of data points from a single class. Higher values are better for all these clustering evaluation metrics.

For the data imputation task, we employed three commonly used evaluation metrics: the Pearson Correlation Coefficient (PCC), L1 loss, and Root Mean Square Error (RMSE). In particular, PCC measures the linear correlation between the imputed and ground truth values, with higher scores suggesting better preservation of global structural patterns in the data. The loss of L1 captures the average absolute deviation between predicted and actual values. Its robustness to outliers makes it a reliable metric for assessing local imputation accuracy. RMSE computes the square root of the mean squared error and emphasizes larger deviations due to its quadratic penalization. A higher PCC reflects better global similarity to the ground truth, while lower L1 loss and RMSE values reflect smaller prediction errors, suggesting more accurate imputations. These metrics provide a comprehensive assessment of both the global consistency and the local precision of the imputed results.

## 4. Discussion

In this study, we present GatorST, a versatile and scalable contrastive meta-learning framework specifically designed for analyzing ST data. GatorST effectively addresses the key challenges in ST data, such as high dimensionality, sparsity, and technical noise, by integrating localized spatial interactions with global gene expression semantics. To capture fine-grained spatial topology, GatorST constructs two-hop neighborhood subgraphs, which are enhanced through cluster-aware pseudo-labeling obtained via k-means clustering. The pseudo-labeling, which serves as weak supervision, improves the contrastive learning process, promoting both spatial coherence and biological relevance in the learned representations. Moreover, GatorST employs an episodic training strategy, enabling adaptive representation learning across various spatial resolutions, gene expression profiles, and tissue architectures.

We validated the effectiveness of GatorST through comprehensive experiments on multiple benchmark datasets, including the human DLPFC, human breast cancer, and mouse brain anterior tissues. The experimental results show that GatorST outperforms state-of-the-art methods in both spatial domain identification and gene expression imputation tasks. These findings highlight GatorST’s potential as a powerful tool for deciphering spatial organization and recovering transcriptomic signals in complex ST data. A detailed parameter sensitivity analysis provides valuable insight into the internal dynamics of GatorST and how different parameter selections impact its performance. The results obtained from the ablation study further demonstrate the efficacy of the GatorST framework. When contrastive learning or subgraph-based modeling was removed, clustering performance decreased significantly. These findings emphasize the critical roles of both contrastive learning and subgraph-based modeling in capturing biological and spatial signals within ST data.

With regard to the research methods, this study has several strengths. (i) The proposed GatorST framework effectively integrates local spatial interactions and global gene expression semantics by modeling localized neighborhoods through graph-based substructures and deriving high-level biological patterns via k-means clustering. This combined contextual representation mitigates spatial biases and improves the biological relevance of the clustering outcomes. (ii) Instead of relying on simplistic corruption strategies or heavy graph data augmentation, GatorST introduces pseudo-label supervision into contrastive learning. This approach strengthens intra-class cohesion and ensures meaningful separation between classes. (iii) The incorporation of an episodic training strategy inspired by meta-learning enables GatorST to adapt to variations in spatial resolution, gene expression profiles, and biological heterogeneity. This adaptability improves generalization across diverse ST datasets. (iv) The joint optimization of contrastive and cross-entropy losses encourages the learning of embeddings that are not only spatially structured but also task-adaptive. This dual-objective approach reduces overfitting and improves the robustness of the model in complex biological contexts.

Although this study has successfully demonstrated the effectiveness of GatorST across multiple benchmarks, its performance on large-scale ST data needs to be systematically investigated. Recent advances in highthroughput spatial transcriptomics technologies have led to the generation of large datasets, which pose significant computational and modeling challenges. The key aspects of dealing with such large-scale data highlight the need for scalable graph construction, memory-efficient continual learning, and maintaining embedding quality across spatial domains with substantial variability in size, density, and topological complexity. Future studies should incorporate advanced techniques such as graph sparsification, hierarchical representation learning, and scalable mini-batch contrastive optimization. These adaptations will be crucial for ensuring the effectiveness of GatorST when handling increasingly large and complex ST datasets.

## Code availability

GatorST is provided as a Python package available at https://github.com/QSong-github/GatorST, with detailed functions for implementation.

## Compliance with Ethics Requirements

This article does not contain any studies with human or animal subjects.

## Declaration of Competing Interest

The authors declare that they have no known competing financial interests or personal relationships that could have appeared to influence the work reported in this paper.

## Acknowledgments

J.B. is supported by National Institutes of Health grants R01AG083039, RF1AG084178, RF1AG077820, R01AG080991, R01AG080624, and R01AG076234.

Q.S. is supported by the National Institute of General Medical Sciences of the National Institutes of Health (R35GM151089).

## Notes

### Competing Interest Statement

The authors have declared no competing interest.

### Summary of Updates

Add a coauthor, Qin Ma, Department of Biomedical Informatics, College of Medicine, The Ohio State University, Columbus, OH, USA;

## References

[1] J. Du, Y.-C. Yang, Z.-J. An, M.-H. Zhang, X.-H. Fu, Z.-F. Huang, Y. Yuan, J. Hou, Advances in spatial transcriptomics and related data analysis strategies, Journal of Translational Medicine 21 (1) (2023) 330.

[2] E. Armingol, A. Officer, O. Harismendy, N. E. Lewis, Deciphering cell– cell interactions and communication from gene expression, Nature Reviews Genetics 22 (2) (2021) 71–88.

[3] S. Jain, M. T. Eadon, Spatial transcriptomics in health and disease, Nature Reviews Nephrology 20 (10) (2024) 659–671.

[4] L. Moses, L. Pachter, Museum of spatial transcriptomics, Nature methods 19 (5) (2022) 534–546.

[5] A. L. Ji, A. J. Rubin, K. Thrane, S. Jiang, D. L. Reynolds, R. M. Meyers, M. G. Guo, B. M. George, A. Mollbrink, J. Bergenstråhle, et al., Multimodal analysis of composition and spatial architecture in human squamous cell carcinoma, Cell 182 (2) (2020) 497–514.

[6] S. G. Rodriques, R. R. Stickels, A. Goeva, C. A. Martin, E. Murray, C. R. Vanderburg, J. Welch, L. M. Chen, F. Chen, E. Z. Macosko, Slide-seq: A scalable technology for measuring genome-wide expression at high spatial resolution, Science 363 (6434) (2019) 1463–1467.

[7] R. R. Stickels, E. Murray, P. Kumar, J. Li, J. L. Marshall, D. J. Di Bella, P. Arlotta, E. Z. Macosko, F. Chen, Highly sensitive spatial transcriptomics at near-cellular resolution with slide-seqv2, Nature biotechnology 39 (3) (2021) 313–319.

[8] A. Chen, S. Liao, M. Cheng, K. Ma, L. Wu, Y. Lai, X. Qiu, J. Yang, J. Xu, S. Hao, et al., Spatiotemporal transcriptomic atlas of mouse organogenesis using dna nanoball-patterned arrays, Cell 185 (10) (2022) 1777–1792.

[9] M. Zhang, S. W. Eichhorn, B. Zingg, Z. Yao, K. Cotter, H. Zeng, H. Dong, X. Zhuang, Spatially resolved cell atlas of the mouse primary motor cortex by merfish, Nature 598 (7879) (2021) 137–143.

[10] S. Shah, Y. Takei, W. Zhou, E. Lubeck, J. Yun, C.-H. L. Eng, N. Koulena, C. Cronin, C. Karp, E. J. Liaw, et al., Dynamics and spatial genomics of the nascent transcriptome by intron seqfish, Cell 174 (2) (2018) 363–376.

[11] S. Codeluppi, L. E. Borm, A. Zeisel, G. La Manno, J. A. van Lunteren, C. I. Svensson, S. Linnarsson, Spatial organization of the somatosensory cortex revealed by osmfish, Nature methods 15 (11) (2018) 932–935.

[12] S. K. Longo, M. G. Guo, A. L. Ji, P. A. Khavari, Integrating single-cell and spatial transcriptomics to elucidate intercellular tissue dynamics, Nature Reviews Genetics 22 (10) (2021) 627–644.

[13] E. Zhao, M. R. Stone, X. Ren, J. Guenthoer, K. S. Smythe, T. Pulliam, S. R. Williams, C. R. Uytingco, S. E. Taylor, P. Nghiem, et al., Spatial transcriptomics at subspot resolution with bayesspace, Nature biotechnology 39 (11) (2021) 1375–1384.

[14] J. Kim, S. Rustam, J. M. Mosquera, S. H. Randell, R. Shaykhiev, A. F. Rendeiro, O. Elemento, Unsupervised discovery of tissue architecture in multiplexed imaging, Nature methods 19 (12) (2022) 1653–1661.

[15] J. Hu, X. Li, K. Coleman, A. Schroeder, N. Ma, D. J. Irwin, E. B. Lee, R. T. Shinohara, M. Li, Spagcn: Integrating gene expression, spatial location and histology to identify spatial domains and spatially variable genes by graph convolutional network, Nature methods 18 (11) (2021) 1342–1351.

[16] H. Ren, B. L. Walker, Z. Cang, Q. Nie, Identifying multicellular spatiotemporal organization of cells with spaceflow, Nature communications 13 (1) (2022) 4076.

[17] V. Singhal, N. Chou, J. Lee, Y. Yue, J. Liu, W. K. Chock, L. Lin, Y.-C. Chang, E. M. L. Teo, J. Aow, et al., Banksy unifies cell typing and tissue domain segmentation for scalable spatial omics data analysis, Nature Genetics 56 (3) (2024) 431–441.

[18] C. Xu, X. Jin, S. Wei, P. Wang, M. Luo, Z. Xu, W. Yang, Y. Cai, L. Xiao, X. Lin, et al., Deepst: identifying spatial domains in spatial transcriptomics by deep learning, Nucleic Acids Research 50 (22) (2022) e131–e131.

[19] J. Li, S. Chen, X. Pan, Y. Yuan, H.-B. Shen, Cell clustering for spatial transcriptomics data with graph neural networks, Nature Computational Science 2 (6) (2022) 399–408.

[20] K. Dong, S. Zhang, Deciphering spatial domains from spatially resolved transcriptomics with an adaptive graph attention auto-encoder, Nature communications 13 (1) (2022) 1739.

[21] B. Wang, J. Luo, Y. Liu, W. Shi, Z. Xiong, C. Shen, Y. Long, Spatial-mgcn: a novel multi-view graph convolutional network for identifying spatial domains with attention mechanism, Briefings in Bioinformatics 24 (5) (2023) bbad262.

[22] Y. Long, K. S. Ang, M. Li, K. L. K. Chong, R. Sethi, C. Zhong, H. Xu, Z. Ong, K. Sachaphibulkij, A. Chen, et al., Spatially informed clustering, integration, and deconvolution of spatial transcriptomics with graphst, Nature Communications 14 (1) (2023) 1155.

[23] H. Xu, H. Fu, Y. Long, K. S. Ang, R. Sethi, K. Chong, M. Li, R. Uddamvathanak, H. K. Lee, J. Ling, et al., Unsupervised spatially embedded deep representation of spatial transcriptomics, Genome Medicine 16 (1) (2024) 12.

[24] R. Lopez, A. Nazaret, M. Langevin, J. Samaran, J. Regier, M. I. Jordan, N. Yosef, A joint model of unpaired data from scrna-seq and spatial transcriptomics for imputing missing gene expression measurements, arXiv preprint 1905.02269 (2019).

[25] T. Biancalani, G. Scalia, L. Buffoni, R. Avasthi, Z. Lu, A. Sanger, N. Tokcan, C. R. Vanderburg, å. Segerstolpe, M. Zhang, et al., Deep learning and alignment of spatially resolved single-cell transcriptomes with tangram, Nature methods 18 (11) (2021) 1352–1362.

[26] L. McInnes, J. Healy, J. Melville, Umap: Uniform manifold approximation and projection for dimension reduction, arXiv preprint 1802.03426 (2018).

[27] F. A. Wolf, F. K. Hamey, M. Plass, J. Solana, J. S. Dahlin, B. Göttgens, N. Rajewsky, L. Simon, F. J. Theis, Paga: graph abstraction reconciles clustering with trajectory inference through a topology preserving map of single cells, Genome biology 20 (2019) 1–9.

[28] I. Korsunsky, N. Millard, J. Fan, K. Slowikowski, F. Zhang, K. Wei, Y. Baglaenko, M. Brenner, P.-r. Loh, S. Raychaudhuri, Fast, sensitive and accurate integration of single-cell data with harmony, Nature methods 16 (12) (2019) 1289–1296.

[29] H. T. N. Tran, K. S. Ang, M. Chevrier, X. Zhang, N. Y. S. Lee, M. Goh, J. Chen, A benchmark of batch-effect correction methods for single-cell rna sequencing data, Genome biology 21 (2020) 1–32.

[30] T. N. Kipf, M. Welling, Semi-supervised classification with graph convolutional networks, arXiv preprint 1609.02907 (2016).

[31] P. Veličković, G. Cucurull, A. Casanova, A. Romero, P. Lio, Y. Bengio, Graph attention networks, arXiv preprint 1710.10903 (2017).

[32] A. Vaswani, N. Shazeer, N. Parmar, J. Uszkoreit, L. Jones, A. N. Gomez, L-.Kaiser, I. Polosukhin, Attention is all you need, Advances in neural information processing systems 30 (2017).

[33] C. Finn, P. Abbeel, S. Levine, Model-agnostic meta-learning for fast adaptation of deep networks, in: International conference on machine learning, PMLR, 2017, pp. 1126–1135.

[34] T. Hospedales, A. Antoniou, P. Micaelli, A. Storkey, Meta-learning in neural networks: A survey, IEEE transactions on pattern analysis and machine intelligence 44 (9) (2021) 5149–5169.

[35] A. Vettoruzzo, M.-R. Bouguelia, J. Vanschoren, T. Rögnvaldsson, K. Santosh, Advances and challenges in meta-learning: A technical review, IEEE transactions on pattern analysis and machine intelligence 46 (7) (2024) 4763–4779.

[36] K. R. Maynard, L. Collado-Torres, L. M. Weber, C. Uytingco, B. K. Barry, S. R. Williams, J. L. Catallini, M. N. Tran, Z. Besich, M. Tippani, et al., Transcriptome-scale spatial gene expression in the human dorsolateral prefrontal cortex, Nature neuroscience 24 (3) (2021) 425– 436.

